# Multiscale fracture resistance of human annulus fibrosus and its deterioration with degeneration

**DOI:** 10.64898/2026.10.20.759657

**Authors:** Xiaoyi Lan, Nathalie R.H. Singh, Tianxiao Xie, Heng Zhu, Kahlan V. Parker, Aram Bahmani, Zhen Yang, Li Li, Yihang Xiao, Hatem M. Titi, Matthew J. Harrington, Lisbet Haglund, Jianyu Li

## Abstract

Soft fibrous tissues must resist fracture under repeated loading, yet how degeneration compromises this resistance remains unclear. We investigated human annulus fibrosus using fracture and fatigue testing, proteoglycan depletion, microscopy, and X-ray scattering. Non-degenerate tissue exhibited pronounced fracture anisotropy, with axial toughness nearly fourfold higher than interlamellar delamination toughness. Degeneration reduced axial and radial toughness despite comparatively preserved tensile modulus and strength. Proteoglycan depletion reduced toughness most strongly in radial and delamination loading. Neither tissue state showed measurable macroscopic crack extension over 20,000 cycles at 70% of its respective monotonic failure strain. Nevertheless, fatigue produced localized structural changes, and degenerate tissue exhibited secondary delamination and diffuse collagen damage. X-ray scattering revealed fatigue- and degeneration-dependent changes in collagen periodicity and intermolecular packing. These findings connect tissue fracture resistance to collagen–matrix interactions across scales and inform the design of crack-resistant biomaterials.

## Introduction

Nature provides many examples of tough materials that achieve crack resistance not through any single component, but through coordinated interactions among structurally distinct components. Nacre, for example, derives its toughness from cooperation between stiff mineral platelets and soft organic interfaces that deflect cracks and dissipate energy. The annulus fibrosus (AF) represents another example. Forming the fibrous outer ring of the intervertebral disc (IVD), the AF contains the nucleus pulposus, redistributes mechanical loads, and resists crack initiation and propagation under complex, repetitive spinal loading over decades of use ^1, 2^. This function progressively deteriorates with disc degeneration, during which small annular fissures can emerge, enlarge, and coalesce into larger defects associated with disc failure and herniation ^3, 4, 5, 6^. Understanding the mechanisms underlying AF crack resistance and its deterioration with degeneration is therefore important because IVD degeneration is a major cause of chronic low back pain and disability worldwide, imposing enormous personal and socioeconomic burdens ^7, 8^.

Crack propagation in AF can occur under a single increasing load or progressively under repetitive subfailure loading, representing distinct mechanical challenges. Monotonic fracture toughness quantifies the energy required to propagate a pre-existing crack during a single loading event ^9, 10, 11^, whereas fatigue resistance reflects the ability of a cracked tissue to withstand repeated cyclic loading without progressive crack extension ^12^. Characterizing these behaviors is further complicated by the hierarchical and strongly anisotropic organization of the AF ^13, 14, 15^, for which mechanical responses depend on tissue orientation, crack path, and structural scale. Conventional tensile properties provide important measures of bulk tissue mechanics, while fracture-mechanics approaches offer complementary information on resistance to crack initiation and propagation. Integrating fracture mechanics with multiscale structural characterization is therefore important for resolving how tissue architecture and matrix composition contribute to crack resistance. Yet how crack resistance emerges from the hierarchical collagen architecture and surrounding matrix, how tissue responds to monotonic versus cyclic loading, and how these responses deteriorate with degeneration remain largely unexplored.

Here, we address these gaps using a multiscale experimental framework integrating direction-dependent monotonic fracture testing, cyclic fatigue loading, selective depletion of chondroitin sulfate-associated matrix components, histological and molecular imaging, and small- and wide-angle X-ray scattering. By comparing non-degenerate and degenerate human AF and resolving responses from tissue-scale fracture to collagen molecular organization, we link crack resistance to collagen architecture, matrix composition, and structural integrity across scales. This approach reveals how collagen and the surrounding matrix jointly contribute to fracture resistance, how repeated subcritical loading generates localized structural alterations before measurable macroscopic crack growth, and how degeneration progressively compromises this multiscale protection. These findings reframe disc degeneration as a progressive, multiscale failure of fracture protection, with direct implications for understanding herniation progression and for the rational design of AF repair biomaterials (Fig. 1 and Supplementary Fig. 1).

**Fig. 1.**
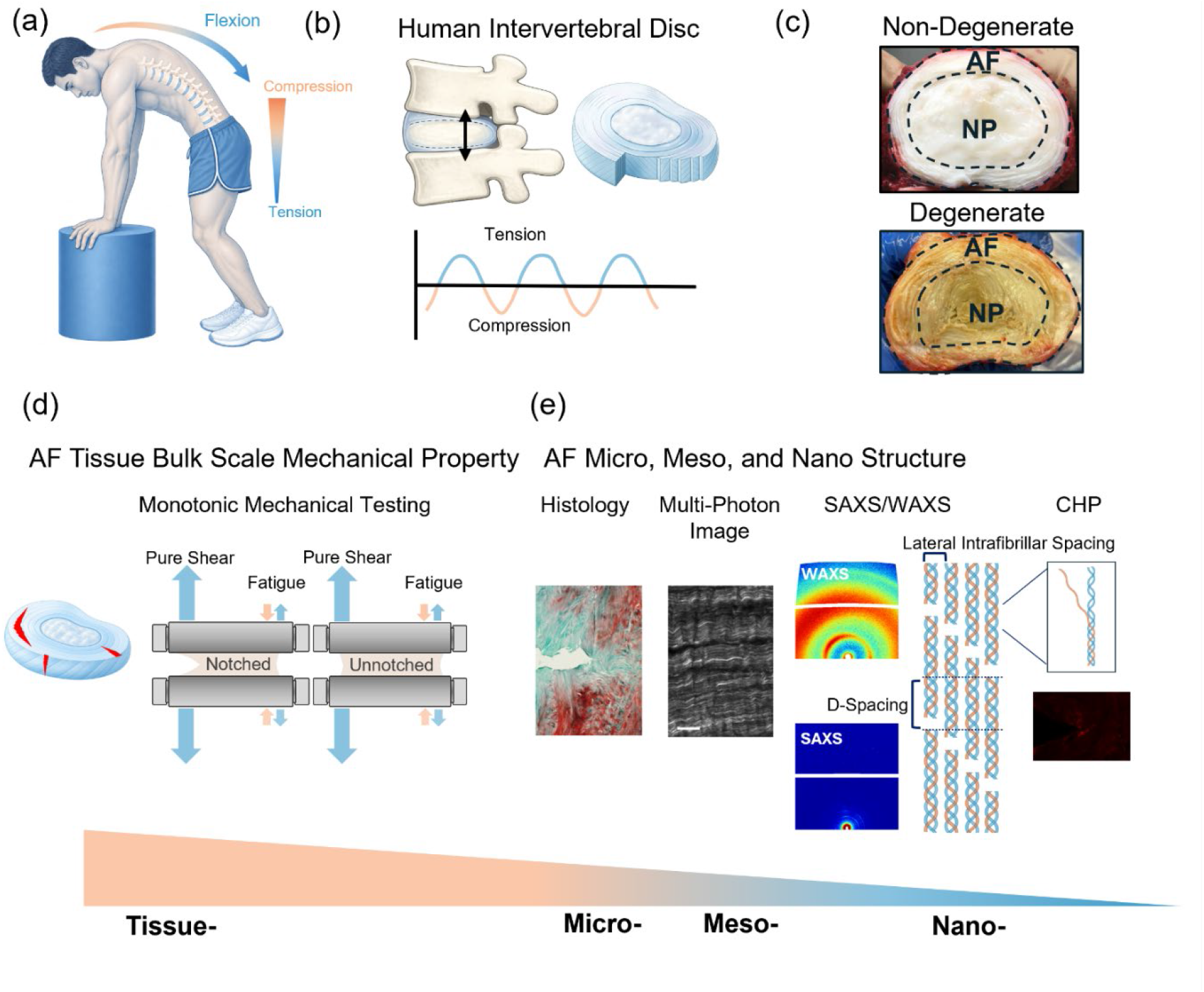
Multiscale framework for investigating fracture behavior of non-degenerate and degenerate annulus fibrosus. a Human spine subjected to repetitive cyclic loading during daily activities. b Intervertebral disc anatomy showing the annulus fibrosus (AF) and nucleus pulposus. c Representative non-degenerate and degenerate human intervertebral discs, highlighting the annulus fibrosus (AF) and nucleus pulposus (NP). d Tissue-scale mechanical testing framework used to characterize AF fracture behavior, including notched and unnotched pure-shear testing under monotonic and cyclic loading. e Multiscale structural characterization of AF tissue spanning the micro-, meso-, and nano-scales using histology, multiphoton imaging, small- and wide-angle X-ray scattering (SAXS/WAXS), and collagen hybridizing peptide (CHP) staining to assess extracellular matrix organization, collagen fiber architecture, fibril periodicity and intermolecular packing, and molecular collagen damage.

## Results

### Fracture toughness of non-degenerate and degenerate AF

We first quantified the resistance of the human AF to crack propagation with fracture toughness, defined as the energy required to create a unit area of crack surface in a material ^11^. Given the crack orientation and dimension of human AF samples, we performed 180° peeling tests for interlamellar delamination (Fig. 2a) and pure shear tests for axial and radial fracture (Fig. 2b,c) ^16, 17^. These tests characterize crack resistance during a single monotonic loading event, in contrast to the cyclic fatigue loading examined subsequently. Human AF exhibits pronounced toughness anisotropy (Fig. 2d). Non-degenerate axial toughness was 8.613 ± 0.406 kJ/m², compared with 6.009 ± 0.722 kJ/m² radially and 2.261 ± 0.816 kJ/m² in delamination. Degenerate specimens had lower corresponding values of 3.944 ± 1.999, 3.005 ± 0.389, and 1.250 ± 0.222 kJ/m². Two-way ANOVA detected an orientation × degeneration interaction (*P* = 0.0208), indicating that degeneration affected fracture toughness differently across loading orientations. Šídák-adjusted comparisons showed significant reductions with degeneration in axial (*P* = 0.0002) and radial (*P* = 0.0072) toughness; the reduction in delamination toughness was not statistically significant. To confirm that the observed directional dependence was not unique to the human specimens, we also tested non-degenerate bovine AF specimens across anatomical orientations, confirming a similar anisotropic trend.

**Fig. 2.**
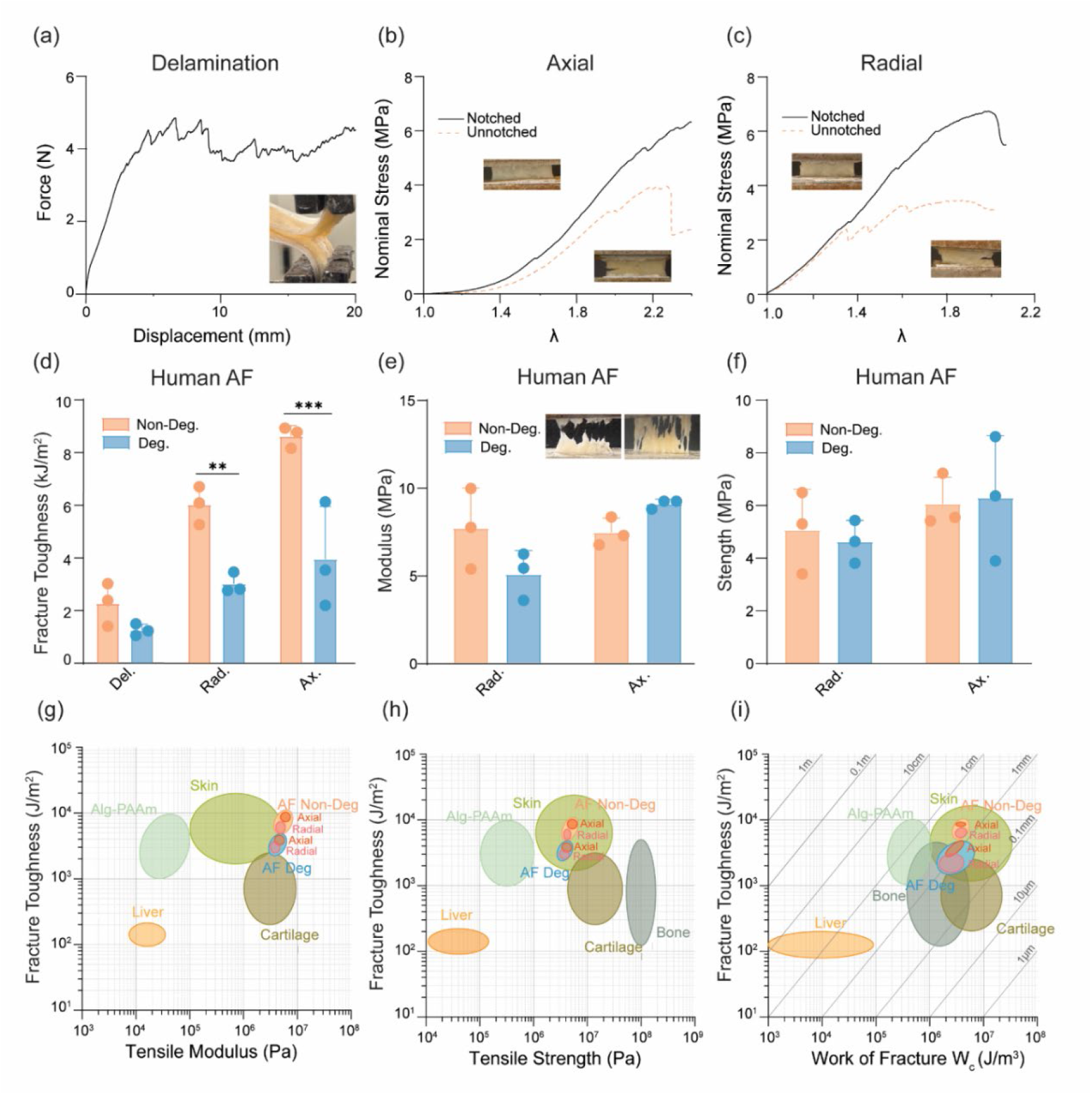
Fracture toughness characterization of human annulus fibrosus across anatomical directions and tissue conditions. a Representative force-displacement curve from delamination testing, with inset showing specimen geometry and crack propagation. b,c Engineering stress-stretch curves for notched and unnotched specimens under axial and radial loading, respectively, with insets showing representative specimens. d Fracture toughness of non-degenerate and degenerate human AF across three anatomical directions (delamination, radial, axial); degeneration significantly reduced toughness in the radial and axial directions. e Tensile modulus of non-degenerate and degenerate human AF in the radial and axial directions, with insets showing representative fracture surface images. f Tensile strength of non-degenerate and degenerate human AF in the radial and axial directions. g–i Ashby plots benchmarking AF fracture toughness against other biological and synthetic tough materials (Alg-PAAm ^19^, skin ^20, 21^, cartilage ^22, 23^, bone ^24, 25, 26^, liver^27, 28^, plotted against g tensile modulus, h tensile strength, and i work of fracture (Wc); diagonal lines are reference contours of characteristic flaw size, used to compare flaw tolerance across material classes. (* *P* < 0.05, ** *P* < 0.01, *** *P* < 0.001, **** *P* < 0.0001; statistical comparisons for panel d by two-way ordinary ANOVA with Šídák’s post-hoc correction for degeneration effects). Data are presented as mean ± SD; n = 3 biological replicates per tissue condition.

Tensile modulus, measured from the initial portion of the stress-stretch curve of unnotched specimens, ranged from 3.56 to 9.93 MPa depending on loading direction (Fig. 2e), but no significant differences were found between non-degenerate and degenerate AF in either axial or radial orientations. Similarly, tensile strength showed no significant degeneration-dependent differences in either loading direction (Fig. 2f). The property maps in Fig. 2g–i contextualize AF fracture toughness, tensile modulus, tensile strength, and work of fracture alongside those of other biological and synthetic materials. In the fracture toughness-tensile strength map, AF exhibits greater fracture toughness than cartilage and liver and values comparable to skin (Fig. 2h). Together, these comparisons illustrate the combination of properties underlying AF mechanical performance, although differences in testing methods and conditions preclude direct comparisons across materials.

### Effect of proteoglycan depletion on AF fracture

Fracture resistance in collagen-rich tissues is often attributed primarily to the collagen network ^18^. However, proteoglycan loss is a defining feature of IVD degeneration. In the AF, proteoglycans are progressively degraded by aggrecanases (ADAMTS-4/5) and metalloproteinases, driven by inflammatory cytokines such as IL-1β and TNF-α, and further amplified by the senescence-associated secretory phenotype. Although these catabolic processes are well established in disc degeneration, whether proteoglycan depletion directly compromises AF fracture resistance and through what interaction with the collagen network still remains unclear.

Gross examination revealed visible differences between non-degenerate and degenerate AF, including yellowing and discoloration of degenerate annular tissue (Fig. 3a). To directly assess the contribution of chondroitin sulfate-rich matrix components to fracture resistance, paired AF specimens were treated with chondroitinase ABC (ChABC) for 24 hours to deplete chondroitin sulfate from the extracellular matrix before mechanical testing (Fig. 3b). Sirius Red/Alcian Blue staining showed abundant proteoglycan-associated staining within non-degenerate AF, whereas degenerate AF exhibited markedly reduced staining across the examined orientations (Fig. 3c,d). ChABC treatment substantially reduced Alcian Blue staining in both tissue states, confirming depletion of chondroitin sulfate-associated matrix components (Fig. 3e,f).

**Fig. 3.**
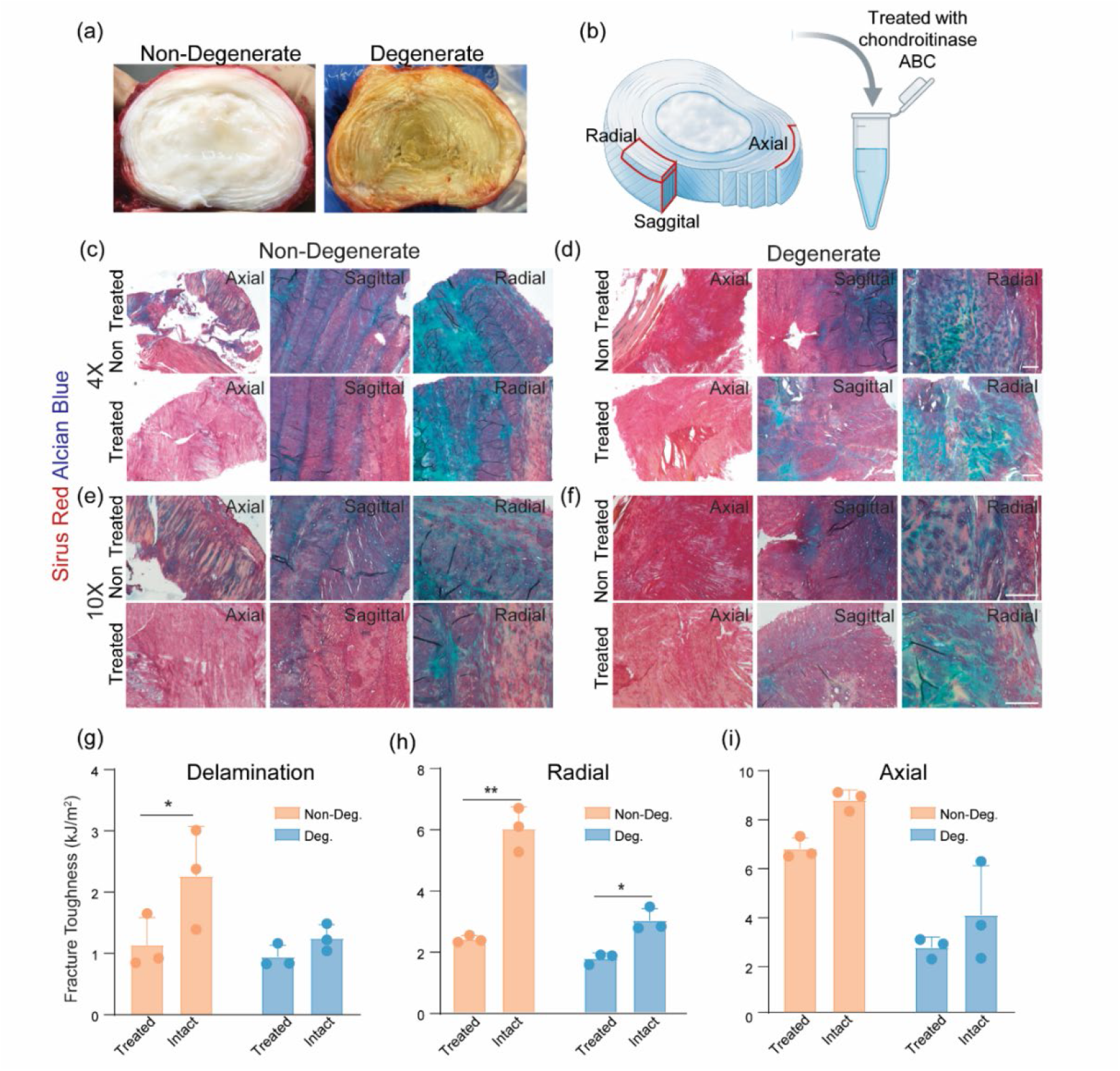
Proteoglycan depletion reduces annulus fibrosus fracture toughness in an orientation- and degeneration-dependent manner. a Representative gross images of non-degenerate and degenerate human annulus fibrosus (AF), showing degeneration-associated tissue discolouration. b Schematic illustrating the axial, radial, and sagittal tissue orientations and enzymatic depletion of chondroitin sulfate-rich proteoglycans using chondroitinase ABC (ChABC). c–f Representative Sirius Red/Alcian Blue-stained sections of non-degenerate c,e and degenerate d,f AF, shown at 4× c,d and 10× e,f magnification, before and after ChABC treatment. Sections are shown in axial, sagittal, and radial orientations. ChABC treatment reduced Alcian Blue staining, consistent with depletion of proteoglycan-associated glycosaminoglycans. g–i Fracture toughness of untreated (intact) and ChABC-treated non-degenerate and degenerate AF in the g delamination, h radial, and i axial directions. Proteoglycan depletion reduced fracture toughness most prominently in the delamination and radial directions, with a smaller effect in the axial direction. Data are shown as individual samples with mean ± SD. (* *P* < 0.05, ** *P* < 0.01, *** *P* < 0.001, **** *P* < 0.0001; statistical comparisons by two-way repeated-measures ANOVA with Šídák’s post-hoc correction for enzymatic treatment effects). Scale bars, 300 µm. n = 3 biological replicates per tissue condition; untreated and treated specimens were matched.

We found that ChABC treatment reduced fracture toughness in an orientation-dependent manner (Fig. 3g–i). In the delamination direction, this treatment significantly reduced fracture toughness in non-degenerate AF, bringing the measured values closer to those of untreated degenerate tissue (Fig. 3g). In the radial direction, ChABC significantly reduced fracture toughness in both non-degenerate and degenerate AF (Fig. 3h). In contrast, the axial direction showed a smaller reduction after ChABC treatment that did not reach statistical significance (Fig. 3i). Quantitatively, relative to untreated controls, the reduction in fracture toughness after ChABC treatment was approximately 56% in the radial direction and 48% for delamination, compared with approximately 24% in the axial direction. Thus, depletion of chondroitin sulfate-associated matrix components had the greatest mechanical effect on radial and interlamellar fracture resistance, which relies on interfibrillar or interlamellar cohesion. Notably, ChABC treatment did not, however, fully reproduce the fracture-toughness phenotype of naturally degenerate AF across all orientations.

Histological analysis provided a structural basis for the directional toughness reduction. Sirius Red/Alcian Blue staining showed that non-degenerate AF contained densely packed, highly aligned collagen fibers within a well-defined lamellar architecture, with abundant proteoglycan-associated staining in the interfibrillar matrix (Fig. 3c,e). Degenerate AF exhibited collagen disorganization, reduced fiber alignment, collagen network disruption, less distinct lamellar boundaries, and reduced Alcian Blue staining. Importantly, degenerate AF also showed more micro-defects and discontinuities (Fig. 3d,f). Thus, proteoglycan depletion weakens AF fracture resistance, and natural degeneration was associated with structural alterations beyond the loss of chondroitin sulfate-rich matrix components alone.

We next examined whether ChABC treatment produced corresponding changes in conventional tensile properties. These effects were more limited and orientation-dependent than those observed for fracture toughness (Supplementary Fig. 2). ChABC treatment significantly reduced radial tensile strength in both non-degenerate and degenerate AF, whereas no significant treatment-dependent difference was observed in axial tensile strength. Tensile modulus decreased significantly after ChABC treatment in non-degenerate radial specimens and degenerate axial specimens, but not in the other groups. By comparison, untreated non-degenerate and degenerate AF did not differ significantly in tensile modulus or tensile strength in either orientation (Fig. 2e,f). Overall, fracture toughness was more sensitive than conventional tensile properties to both degeneration and ChABC treatment, identifying fracture resistance as a mechanical phenotype that more readily captures degeneration-associated changes in matrix support and collagen architecture.

### Microstructure of fatigued AF tissue

While monotonic fracture testing characterises crack resistance under a single large loading, physiological loading of the spine is inherently cyclic with millions of cycles of lower loading over a lifetime. To examine crack behaviour under repeated sub-failure loading, we performed cyclic fatigue tests. With a customized fatigue testing machine, we loaded pre-notched human AF specimens cyclically to 70% of their respective monotonic failure strain for up to 20,000 cycles (Fig. 4a). Interestingly, no measurable macroscopic crack extension was detected in either non-degenerate or degenerate AF during the loading protocol (Fig. 4b–c). Despite the absence of measurable crack propagation, the crack-tip morphology differed markedly between degeneration states (Fig. 4d). Non-degenerate AF retained a relatively sharp and well-defined crack tip in both radial and axial specimens. In contrast, degenerate AF exhibited a broader and more distorted crack-tip region, with additional fissures extending preferentially along the delamination direction. Bovine AF similarly tolerated repeated sub-failure loading without obvious macroscopic crack propagation, while exhibiting pronounced cyclic hysteresis, indicating substantial mechanical energy dissipation during repeated loading (Supplementary Fig. 3).

**Fig. 4.**
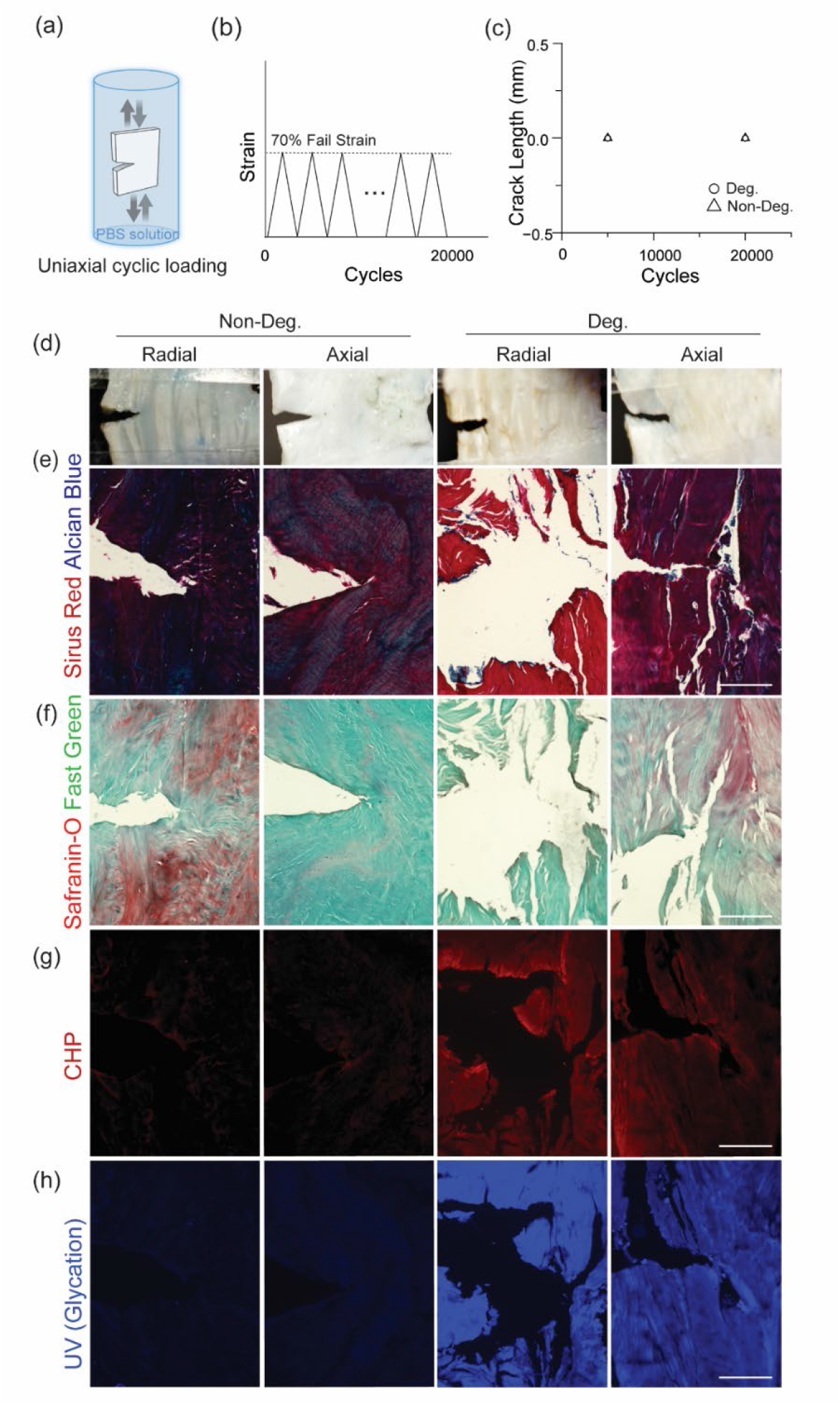
Degeneration alters the structural and molecular response of annulus fibrosus to cyclic fatigue loading. a Schematic of uniaxial cyclic loading of notched annulus fibrosus (AF) specimens immersed in PBS. b Specimens were cyclically loaded to 70% of their monotonic failure strain for 20,000 cycles. c Crack extension as a function of cycle number, showing no measurable macroscopic crack propagation during fatigue loading. d Representative brightfield images of radial and axial specimens from non-degenerate and degenerate AF following cyclic loading. e,f Sirius Red/Alcian Blue and Safranin-O/Fast Green staining showing collagen organisation and proteoglycan distribution around the crack tip. g Collagen hybridising peptide (CHP) staining showing collagen denaturation associated with fatigue-induced matrix damage. h UV autofluorescence imaging showing glycation-associated fluorescence. Despite the absence of measurable crack extension, non-degenerate and degenerate AF exhibited distinct crack-tip morphologies and matrix-level responses, indicating that degeneration alters the mechanisms by which AF accommodates cyclic loading. Scale bars, 300 µm.

Histological analysis revealed distinct spatial patterns of matrix alteration around the crack tip after fatigue loading. Sirius Red/Alcian Blue staining showed densely packed and aligned collagen fibers converging toward the crack tip in non-degenerate AF, with Alcian Blue staining retained in the surrounding matrix (Fig. 4e). Degenerate-fatigued AF exhibited disrupted collagen organization and markedly reduced Alcian Blue staining around the crack region. Safranin-O/Fast Green staining further showed a localized loss of Safranin-O signal at the crack tip in non-degenerate AF despite preservation in surrounding regions, whereas degenerate AF showed broadly reduced Safranin-O staining without a clear crack-tip-specific pattern (Fig. 4f).

Collagen hybridizing peptide (CHP) staining further revealed corresponding differences in collagen molecular integrity. In non-degenerate AF, CHP staining was location-dependent within the fatigue-tested specimen: it was concentrated at the crack tip, with minimal staining in the intact tissue away from the crack tip. In degenerate AF, CHP staining was present throughout the specimen regardless of location relative to the crack tip. UV autofluorescence imaging additionally revealed strong, diffuse glycation-associated fluorescence, consistent with the accumulation of advanced glycation end-products (AGEs), in degenerate AF, compared with minimal signal in non-degenerate tissue (Fig. 4h). Deep-tissue multiphoton images of intact and fatigued bovine AF are presented in Supplementary Fig. 4.

### X-ray scattering assessments

To further determine whether cyclic loading altered collagen structure in nanoscale, we used X-ray scattering techniques to quantify collagen fibril periodicity and intermolecular packing of AF. We characterized both radial and axial cuts of non-degenerate and degenerate IVD specimens before and after fatigue loading. Because radial cuts span multiple lamellae, they capture a greater contribution from the interlamellar and proteoglycan-associated matrix than axial cuts, which sample fewer lamellae. In each sample, three locations along the crack path were analyzed after fatigue: the crack-tip region immediately adjacent to the crack tip, the mid-crack region approximately halfway along the crack path, and the crack-end region within the terminal one-sixth of the crack path. Regional measurements are presented in Supplementary Fig. 5.

SAXS measurements probe the d-periodicity originating from the quarter stagger of triple helices within assembled collagen fibrils (Fig. 5b,c). In radial specimens, the d-spacing of intact tissue was 64.61 ± 0.30 nm in non-degenerate and 64.42 ± 0.65 nm in degenerate specimens, consistent with the established ∼64 nm d-period of dried type I collagen ^29, 30^. Two-way repeated-measures (RM) ANOVA revealed a significant effect of fatigue (*P* = 0.0310), but no significant difference between degenerate and non-degenerate samples under a given fatigue condition. After fatigue, at the crack tip, d-spacing decreased to 62.81 ± 0.62 nm in non-degenerate AF and 61.87 ± 0.89 nm in degenerate AF. Pairwise comparisons between intact and crack-tip measurements within each degeneration group did not show a significant difference in d-spacing. In axial specimens, intact-tissue d-spacing was 64.22 ± 0.09 nm in non-degenerate AF and 64.27 ± 0.10 nm in degenerate AF. Two-way RM ANOVA revealed significant main effects of both fatigue (*P* = 0.0137) and degeneration (*P* = 0.0085), with no significant fatigue × degeneration interaction, providing no evidence that the effect of fatigue differed between non-degenerate and degenerate AF. Pairwise comparisons showed that d-spacing was significantly reduced at the crack tip relative to intact tissue in the degenerate AF group, whereas the corresponding difference in non-degenerate AF was not significant.

**Fig. 5.**
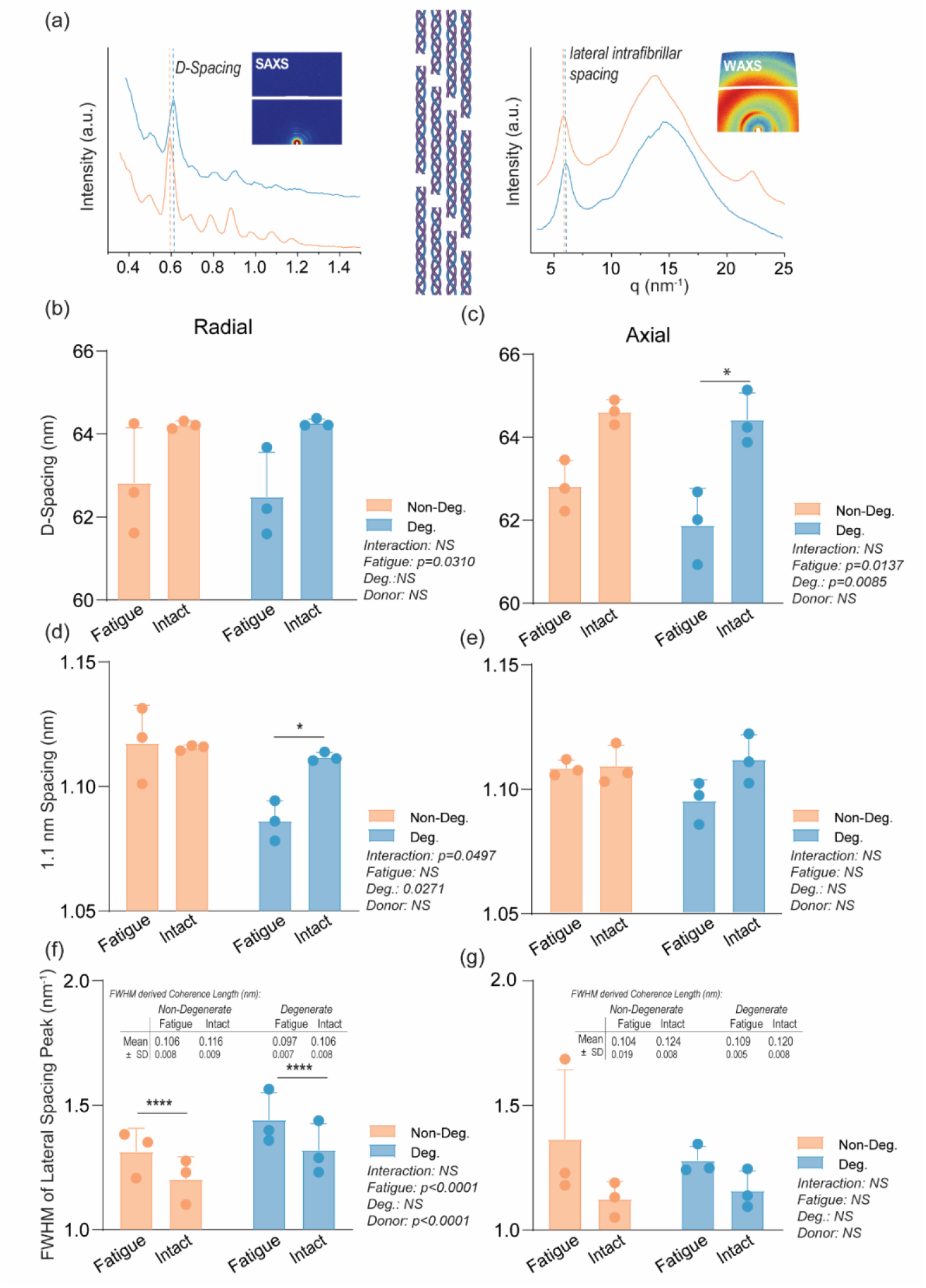
Multiscale collagen structural characterisation of AF tissue using SAXS and WAXS. a Schematic illustration of the hierarchical collagen organisation in AF tissue, from fiber bundles to individual fibrils, with representative 2D SAXS and WAXS scattering patterns. SAXS was used to assess fibril-level structure via the axial D-spacing, while WAXS was used to evaluate molecular-level organisation via the lateral intrafibrillar spacing and its coherence length. b,c Axial D-spacing derived from SAXS in radial and axial specimens, respectively, reflecting the characteristic ∼64 nm periodicity of dried type I collagen fibrils. d,e Lateral intrafibrillar spacing (∼1.1 nm) derived from WAXS in radial and axial specimens, respectively, reflecting collagen intermolecular packing density. f,g Full width at half maximum (FWHM; nm-1) of the 1.1 nm WAXS peak in radial and axial specimens, respectively, as an index of fibril organisational regularity. (* *P* < 0.05, ** *P* < 0.01, *** *P* < 0.001, **** *P* < 0.0001; statistical comparisons by two-way repeated-measures ANOVA with Šídák’s post-hoc correction for fatigue effects). Data are presented as mean ± SD; n = 3 biological replicates per tissue condition.

WAXS measurements further revealed degeneration-dependent changes in lateral intermolecular helix to helix spacing (Fig. 5d,e). In radial specimens, intact lateral spacing was 1.116 ± 0.001 nm in non-degenerate AF and 1.112 ± 0.002 nm in degenerate AF, consistent with the approximately 1.1-nm lateral intermolecular spacing of dried type I collagen ^29^. Two-way RM ANOVA revealed a significant fatigue × degeneration interaction (*P* = 0.0497), indicating that the effect of fatigue on lateral intermolecular spacing differed between non-degenerate and degenerate AF. Neither fatigue nor degeneration showed a significant main effect. Pairwise comparisons revealed that lateral spacing was significantly reduced following fatigue in degenerate specimens (1.086 ± 0.008 nm at the crack tip), while no significant change was observed in non-degenerate specimens following fatigue (1.117 ± 0.05 nm at the crack tip). This interaction suggests that fatigue-induced lateral fibril compaction is specific to proteoglycan-depleted tissue. In axial specimens, no significant effects of fatigue, degeneration, or their interaction were detected for lateral intrafibrillar spacing, though borderline trends were observed for both fatigue (*P* = 0.0622) and the fatigue × degeneration interaction (*P* = 0.0834). These trends mirror the pattern observed in radial specimens, suggesting that a qualitatively similar fatigue- and degeneration-dependent response may be present in axial tissue, but below the threshold of detection at the current sample size.

Lastly, the full width at half maximum (FWHM) of the d ∼1.1-nm WAXS peak was analyzed with Scherrer analysis to determine coherence length, providing an estimate of the spatial ordering of this recurring lateral spacing within collagen fibrils ^31^. In radial specimens, two-way RM ANOVA revealed a highly significant effect of fatigue on the FWHM of the 1.1 nm lateral spacing peak (*P* < 0.0001), with no significant effect of degeneration or fatigue × degeneration interaction, indicating that cyclic loading altered collagen intermolecular fibril packing/coherence. Crack-tip specimens exhibited markedly higher FWHM values (lower coherence lengths) than intact specimens in both non-degenerate (1.314 ± 0.093 vs. 1.203 ± 0.091 nm^-1^) and degenerate (1.441 ± 0.109 vs. 1.319 ± 0.107 nm^-1^) groups, indicating a fatigue-induced reduction in the coherence length within the lateral packing of fibrils. Pairwise comparisons confirmed significant differences between crack-tip and intact specimens in both non-degenerate and degenerate groups (*P* < 0.0001). In axial specimens, no significant effects were detected for any source of variation (all *P* > 0.14), in contrast to the fatigue-dependent changes observed in radial specimens.

## Discussion

The present study shows that fracture resistance in the human annulus fibrosus (AF) arises from coordinated contributions of hierarchical collagen architecture and the surrounding matrix. AF exhibited pronounced fracture anisotropy, with the greatest toughness in the axial direction, intermediate toughness in the radial direction, and the lowest toughness during interlamellar delamination. These directional differences are reported with other fibrous biological tissues, where crack propagation resistance scales with the degree of fiber engagement along the crack path ^20, 32, 33^. Degeneration substantially reduced both axial and radial fracture toughness, indicating a deterioration of the mechanisms that resist crack propagation as the tissue degenerates.

The high axial fracture toughness of non-degenerate AF is consistent with its angle-ply lamellar architecture. Axial crack propagation requires substantial engagement and rupture of collagen fibers oriented approximately ±30° relative to the spinal axis ^34, 35^, whereas radial and interlamellar cracks can advance along pathways that depend more strongly on interfibrillar and interlamellar cohesion. The particularly low delamination toughness therefore highlights the mechanical vulnerability of the interfaces between neighboring lamellae and is consistent with the frequent occurrence of concentric fissures in degenerate discs ^36, 37, 38, 39^. These findings indicate that AF toughness depends not only on collagen fiber strength, but also on how collagen architecture and matrix cohesion interact along different crack paths.

Degenerate AF simultaneously exhibited lower fracture toughness under monotonic loading, but without producing corresponding reductions in tensile modulus or strength. The reduced toughness is related to collagen disorganization, disrupted lamellar boundaries, and increased structural defects. Similarly, the preservation of stiffness and strength likely reflects the competing mechanical consequences of degeneration. Proteoglycan loss and disruption of the interfibrillar matrix can reduce hydration, matrix cohesion, and load transfer. At the same time, tissue dehydration and an increased relative contribution of collagen to the solid matrix can increase apparent stiffness, while glycation-associated collagen crosslinking may further increase resistance to bulk deformation. Strong CHP signal in degenerate specimens indicates that collagen molecular damage was already present before experimental loading, consistent with previous reports of elevated CHP staining in degenerate disc tissue compared with non-degenerate controls ^40^. UV autofluorescence confirmed diffuse AGE accumulation in degenerate AF compared with minimal signal in non-degenerate controls. Because AGEs form non-enzymatic collagen crosslinks that stiffen the fibrillar network and restrict fibril mobility ^41, 42, 43^, their accumulation may impair energy dissipation mechanisms that depend on collagen recruitment, fibril sliding, and molecular unfolding ^43, 44, 45, 46^. Conventional tensile properties therefore reflect the net effect of several potentially opposing structural changes.

Fracture toughness probes a different aspect of tissue mechanics. Whereas tensile modulus and strength characterize the bulk response of relatively intact tissue under distributed loading, fracture toughness reflects the ability of the tissue to redistribute highly concentrated stresses around an existing defect. This resistance to crack propagation depends on mechanisms including fiber recruitment and bridging, interfibrillar sliding, matrix-mediated load transfer, crack deflection, and interlamellar cohesion. Degeneration can compromise these mechanisms even when compensating compositional changes preserve bulk stiffness or strength. Consequently, this distinction suggests that conventional mechanical testing may underestimate degeneration-associated deterioration in damage tolerance. A tissue can remain relatively stiff under homogeneous loading while becoming increasingly vulnerable to propagation of a pre-existing crack. In the AF, where microscopic fissures can accumulate over time, fracture mechanics may therefore provide a particularly relevant measure of progressive structural deterioration.

The proteoglycan-rich matrix contributes to fracture resistance through several complementary mechanisms, including maintaining tissue hydration and osmotic swelling pressure that pre-strain the collagen network ^47, 48, 49^, mediating viscoelastic energy dissipation under cyclic loading ^50^, and providing interfibrillar cohesion that resists lamellar separation ^51^. With degeneration, proteoglycan depletion and associated matrix changes can impair these functions.

Selective proteoglycan depletion provides direct evidence, and further suggests that this matrix contribution is strongly dependent on crack direction. ChABC treatment produced the largest reductions in radial and delamination toughness, whereas the reduction in axial toughness was smaller. This directional response is consistent with the different structural mechanisms governing crack propagation. During delamination, cracks propagate primarily along interfaces between adjacent lamellae, where proteoglycans and other non-collagenous matrix components contribute substantially to interlamellar cohesion ^52^. Accordingly, the pronounced reduction in delamination toughness after ChABC treatment indicates an important role of the interlamellar matrix in resisting lamellar separation. Radial crack propagation similarly requires coordinated deformation and separation among collagen fibers, interfibrillar matrix, and successive lamellae, making it particularly sensitive to loss of matrix-mediated cohesion and load transfer. In contrast, axial fracture requires greater direct engagement and rupture of collagen fibers and therefore depends more strongly on the collagen architecture itself.

ChABC treatment only partially reproduced the reduction in toughness observed with degeneration, indicating that proteoglycan loss is important but not sufficient to explain the full mechanical phenotype. Histology showed reduced proteoglycan staining after enzyme treatment and in degenerate AF; however, because proteoglycan content was not quantified, the extent of depletion cannot be assumed to be equivalent between groups. In addition, chondroitinase ABC primarily cleaves chondroitin sulfate and dermatan sulfate chains ^53^, while other matrix components that contribute to hydration and matrix organization, such as hyaluronan, may remain at least partly preserved. Together, these findings show that AF toughness arises from direction-dependent cooperation between the collagen architecture and surrounding matrix, both of which are progressively compromised by degeneration.

Cyclic loading at 70% of the monotonic failure strain for 20,000 cycles produced no measurable macroscopic crack extension in either non-degenerate or degenerate human AF. Thus, both tissue states retained substantial resistance against fatigue crack under the testing conditions. This finding is consistent with prior studies showing that sub-failure cyclic loading of isolated AF does not necessarily compromise ultimate mechanical properties ^12, 54^, and in other collagen tissue where the crack is arrested during repeated loading ^50^. The pronounced hysteresis observed during cyclic loading further indicates substantial energy dissipation, while multiphoton imaging showed collagen fibers transitioning from a crimped to a straightened configuration, consistent with progressive fiber recruitment and redistribution of load before fiber failure. Similar findings have been reported in other collagenous tissues, where collagen triple-helical denaturation can accumulate progressively with increasing fatigue cycles and creep strain ^55^.

Importantly, the absence of macroscopic crack growth did not imply an absence of localized crack-tip response. While the crack tip of non-degenerate AF remained relatively sharp after cyclic loading, substantial structural change is observed. Histological and molecular analyses revealed a spatially confined response surrounding the crack. Safranin-O staining was reduced locally at the crack tip while remaining evident in the surrounding matrix, and CHP staining was restricted to the crack-tip region in fatigued specimens but absent from intact controls. These findings indicate localized matrix alteration and collagen molecular unfolding rather than diffuse bulk damage, consistent with prior observations that sub-failure fatigue does not produce widespread collagen denaturation in healthy AF ^54^.

Degenerate AF exhibited a different crack-tip response. Under the same normalized loading protocol, crack tips became more distorted and secondary damage developed along the delamination direction. This is consistent with the lower monotonic fracture toughness of degenerate AF and suggests a lower threshold for local structural deformation. Diffuse CHP staining in degenerate tissue also indicates pre-existing collagen molecular damage, which may reduce the additional deformation required to activate further local damage. The secondary delamination further suggests that weakened interlamellar cohesion provides a preferential pathway once crack-tip deformation develops.

Nevertheless, degenerate AF also remained resistant to measurable fatigue crack extension over 20,000 cycles. This should be interpreted in the context of the loading protocol: each group was cycled at 70% of its own monotonic failure strain, so the degenerate specimens experienced a lower absolute loading level consistent with their reduced fracture toughness. In addition, crack-tip blunting may still deconcentrate the stress at crack tip ^56^. In fibrous soft tissues, collagen reorientation, densification, bridging, and strain hardening can blunt the crack tip and reduce the local driving force for crack advance ^18, 57^. This reorientation is driven by fibril rotation and recruitment into alignment with the loading axis, dissipating energy through fibril stretching and sliding ^58^. In degenerate AF, however, the accompanying crack-tip distortion and secondary delamination suggest that this response is less controlled and occurs in a structurally compromised tissue.

X-ray scattering enabled analysis of the effect of fatigue response on collagen fibril periodicity and intermolecular packing. Intact AF exhibited a collagen d-period of 64.479 ± 0.173 nm across degeneration states and specimen orientations, consistent with previous reports ^30^. Thus, the marked differences in tissue-level fracture behavior were not accompanied by gross disruption of baseline collagen periodicity. Following fatigue, however, d-period decreased to 62.501 ± 0.445 nm in crack-associated regions, corresponding to a residual reduction of approximately 2.9 ± 0.7%. This reduction was more pronounced in axial than radial specimens, further suggesting that the nanoscale fatigue response depends on tissue architecture. Because these measurements were obtained after unloading rather than *in situ*, the reduced d-period represents a residual structural state rather than the instantaneous fibril deformation during loading.

One possible interpretation is partial relaxation of the tensile pre-strain carried by collagen fibrils in intact AF ^47^. The AF is not stress free-even in the absence of external loading. Michalek et al. demonstrated substantial residual circumferential strains across the AF, with residual compression in the inner AF and tension toward the outer periphery, and proposed that regional differences in proteoglycan content may contribute to this residual stress state ^47^. They further suggested that proteoglycan loss with degeneration could diminish these beneficial residual stresses ^47^. At the fibrillar scale, a related mechanism has been demonstrated in articular cartilage, where proteoglycan-driven osmotic swelling maintains collagen fibrils under approximately 2% tensile pre-strain, as estimated from collagen d-period measurements. During stress relaxation following compression, the collagen d-period transiently decreased after peak load, indicating relaxation of fibrillar tensile pre-strain and increased intrafibrillar disorder ^59^. These findings link collagen d-period to fibrillar pre-strain and to mechanical coupling between collagen and the proteoglycan-water matrix. Although cartilage differs from AF in composition and loading mode, together these studies provide a mechanistic basis for interpreting the post-fatigue reduction in d-period observed here as a residual alteration in collagen-matrix mechanical coupling. In contrast, chemically fixed bovine pericardial membranes showed little change in collagen d-period after 30,000 loading cycles despite substantial fiber de-crimping and reorientation^50^. Chemical fixation performed before loading may constrain fibril-level rearrangement and alter collagen-matrix interactions during and after cyclic loading. Together, this comparison suggests that changes in d-period under repeated loading may depend strongly on the mechanical coupling between collagen fibrils and their surrounding matrix.

Lateral intermolecular packing showed an additional degeneration-dependent response. A significant degeneration × fatigue interaction indicated that the effect of fatigue depended on the degeneration status: fatigue reduced the approximately 1.1 nm intermolecular spacing in degenerate AF whereas no reduction was observed in non-degenerate AF. Because hydration strongly influences lateral collagen packing, as intermolecular spacing can decrease from approximately 1.5 nm in hydrated collagen to approximately 1.1 nm following dehydration ^29^, all specimens were dehydrated under identical conditions before analysis. The observed differences are therefore unlikely to reflect differences in hydration state at the time of measurement. Instead, the observed differences may reflect a residual structural consequence of fatigue experienced while the tissue was hydrated. Loss of proteoglycans and associated osmotic swelling pressure in degenerate AF may reduce lateral support of the collagen network during cyclic loading due to lowered water retention, allowing greater intermolecular compaction that persists after unloading and standardized dehydration ^52, 60, 61^. This interpretation is consistent with the established role of the proteoglycan-water matrix in collagen-matrix mechanical coupling and in regulating fibrillar spacing and deformation ^29^.

Fatigue also significantly increased the FWHM of the approximately 1.1-nm WAXS peak in radial specimens. Because the width of this equatorial reflection reflects the degree of lateral periodic order in collagen molecular packing, peak broadening is consistent with increased heterogeneity in intermolecular spacing and, via Scherrer analysis, a reduced lateral coherence length of the triple helical packing ^62^. Radial specimens traverse multiple lamellae containing collagen fibrils arranged at alternating angles relative to the spinal axis and therefore sample a more complex multilamellar architecture than axial specimens ^63^. This architecture may make radial specimens particularly susceptible to heterogeneous nanoscale rearrangement during cyclic loading. It should be noted that differential straining on the tissue could additionally impact the FWHM of this peak (and thus the coherence length); however, it is difficult to denote this role within the tissue without excising and measuring single lamella. Notably, the FWHM increase was observed in both non-degenerate and degenerate AF, indicating that this fatigue-induced alteration in collagen packing is distinct from the degeneration-dependent reduction in intermolecular spacing described above. Together, these findings suggest two complementary nanoscale responses to fatigue: a degeneration-dependent lateral compaction of collagen molecules and an architecture-dependent increase in the heterogeneity, or reduced coherence, of their lateral packing.

Several limitations define the scope of this study. First, the cyclic-loading experiments were performed at a single normalized loading level, 70% of the monotonic failure strain, for 20,000 cycles without observing steady macroscopic crack extension. They therefore establish resistance to crack growth under a defined subcritical condition but do not determine a fatigue threshold or endurance limit. Because each group was loaded relative to its own failure strain, equivalent normalized strains also do not necessarily correspond to equivalent stresses or crack-driving forces across degeneration states. Second, the nanoscale measurements were performed after fatigue rather than *in situ*. Accordingly, the observed changes in collagen d-period and intermolecular packing represent persistent post-loading structural states rather than the temporal evolution of fibril deformation during loading. Future *in situ* SAXS/WAXS measurements under hydrated cyclic loading could resolve the reversibility and sequence of these nanoscale changes in real time. Third, natural degeneration also involves concurrent changes in proteoglycan content, hydration, collagen organization, crosslinking, and tissue microstructure. Chondroitinase ABC treatment therefore isolates one component of this process rather than reproducing the full degenerative phenotype. Further investigations that independently manipulate matrix composition, hydration, collagen crosslinking, and fiber architecture could help distinguish their individual and combined contributions to AF fracture. Similarly, donor age, anatomical location, and inter-individual variability may contribute to the mechanical and structural differences observed between degeneration groups and should be addressed in larger donor-resolved studies.

In conclusion, fracture and fatigue resistance in the human AF emerges from complementary contributions of hierarchical collagen architecture and the surrounding matrix. This organization produces pronounced directional toughness, while degeneration selectively compromises resistance to crack propagation even when conventional tensile properties remain comparatively preserved. Proteoglycan-associated matrix components contribute particularly strongly to radial and interlamellar fracture resistance, whereas cyclic loading reveals molecular and fibrillar damage that can accumulate without measurable macroscopic crack extension. Across scales, these findings indicate that non-degenerate AF dissipates and redistributes mechanical energy through collagen realignment, matrix-mediated load transfer, fibrillar rearrangement, and localized molecular damage, thereby limiting crack propagation. Degeneration progressively disrupts this coordinated response through loss of matrix support, collagen disorganization, pre-existing molecular damage, and altered fibril-level mechanics.

These findings have implications for both assessment and repair of disc degeneration. Fracture toughness and fatigue-induced structural changes may reveal loss of damage tolerance that is not apparent from conventional measurements of stiffness or strength alone. Likewise, effective AF repair materials should reproduce not only the bulk mechanical properties of native tissue but also its hierarchical fiber architecture, matrix-mediated load transfer, and capacity to localize and dissipate damage around defects. More broadly, the results establish the AF as a multiscale tough biological composite and provide design principles for engineering soft fibrous materials that must resist repeated loading and crack propagation.

## Methods

### Tissue sample preparation

Human AF specimens were obtained from six organ donors (five male and one female; age range, 21– 72 years) from the L3–L4 and L4–L5 intervertebral discs. Disc degeneration was assessed macroscopically using the Thompson grading system. Donors with Thompson grade I discs were classified as non-degenerate, while donors with Thompson grades III–V were classified as degenerate. Donor demographics, disc levels, and individual Thompson grades are summarized in Supplementary Table S1. Bovine caudal discs (C1-C2 to C3-C5 levels) were obtained frozen from a local butcher. Human tissues were collected from consenting organ donors under institutionally approved procedures and processed following approval from the McGill University Research Ethics Board (IRB #2019-4896 and A03-M11-21A).

AF specimens were harvested from the anterior region of the disc and prepared in radial, axial, and delamination fracture orientations (Supplementary Fig. 1). Single-edge notches were introduced to generate the corresponding crack propagation directions ^37, 64^. Radial specimens were sectioned transversely and notched from the outer AF to promote crack propagation radially inward across successive lamellae. Axial specimens were sectioned circumferentially as ∼2-mm-thick outer-AF strips, with cracks propagating parallel to the spinal axis. Delamination specimens were likewise sectioned circumferentially, but the notch was positioned along an interlamellar interface to promote separation between adjacent lamellae. Thus, the radial and axial configurations probed translamellar fracture along orthogonal directions, whereas the delamination configuration probed interlamellar fracture.

### Fracture testing

Fracture toughness of bovine and human AF specimens was measured using pure-shear testing on an Instron universal testing machine (5965B17167). Intact and single-edge-notched specimens with radial or axial crack orientations were loaded to failure ^10, 65^. Human disc samples were stored at −80 °C after biobank collection, while bovine caudal discs were obtained freshly from a local butcher and subsequently stored at −80 °C; all specimens underwent only a single freeze-thaw cycle before testing.

After thawing, specimens were sectioned to the required geometry using a razor blade, equilibrated overnight at 4 °C in phosphate-buffered saline (PBS), and mechanically tested within 24 h of thawing. Single-edge notches were then introduced using a scalpel blade according to the designated radial or axial crack orientation. To remove slack and minimize handling-induced prestress, specimens were preloaded to 0.05 MPa and preconditioned for 10 cycles to 5% strain. Fracture toughness in the delamination direction was measured separately using a 180° peel test to characterize interlamellar fracture resistance ^66, 67, 68^.

### Cyclic fatigue test

Cyclic fatigue tests were performed using a custom-built fatigue tester equipped with a NEMA 19 stepper motor (Supplementary Fig. 6). To examine fatigue crack behavior across degeneration states, edge-notched human AF specimens from non-degenerate and degenerate discs were subjected to displacement-controlled cyclic loading at 70% of their respective failure strain for 20,000 cycles at 1 Hz. All tests were conducted in PBS bath at room temperature.

For bovine AF, fatigue testing was performed using an Instron universal testing machine (5965B17167). Specimens harvested from frozen bovine caudal discs were prepared in axial and radial orientations and equilibrated overnight in PBS before testing. Edge-notched specimens were subjected to uniaxial displacement-controlled cyclic loading at either 50% or 70% of the failure strain, estimated from adjacent unnotched tissue, for 5,000 or 20,000 cycles at 0.25 Hz. All bovine tests were conducted in a PBS bath at room temperature. Hysteresis loops were generated by plotting stress as a function of strain for individual loading-unloading cycles. Loop areas were determined by numerical integration of the loading and unloading curves.

### Chondroitinase ABC treatment

To assess the contribution of proteoglycans to AF fracture behavior, specimens were treated with chondroitinase ABC (Sigma Aldrich, C3667) before mechanical testing. Chondroitinase ABC was prepared at a final activity of 1.0 U/mL in a pH 8.0 digestion buffer containing 100 mM sodium acetate and 100 mM Tris-HCl. The enzyme stock was prepared at 30 U/mL in 0.01% BSA solution. Specimens were fully submerged in the digestion solution and incubated at 37 °C for 24 h at a solution-to-tissue ratio of 1 mL/g before mechanical testing. Matched untreated control specimens were incubated in the same digestion buffer without chondroitinase ABC at 37 °C for 24 h, using the same solution-to-tissue ratio, before mechanical testing.

### Histology

Following fatigue testing, crack-tip regions from post-fatigue and intact specimens were immediately fixed in 4% paraformaldehyde (PFA), processed, paraffin embedded, and sectioned at 5 µm thickness for structural and molecular characterization. Safranin-O/Fast Green staining was used to assess proteoglycan distribution, whereas Sirius Red/Alcian Blue staining was used to evaluate collagen organization and matrix composition, following standard protocols.

### Imaging of collagen damage and AGE accumulation

Collagen damage at the crack tip was assessed using collagen hybridizing peptide staining (CHP-R; 3Helix, Salt Lake City, UT), and advanced glycation end-product (AGE) accumulation was evaluated by UV-induced tissue autofluorescence using EVOS fluorescence microscopy. For CHP staining, CHP-R solution (20 µM) was heated to 80 °C for 5 min to dissociate the trimeric peptide structure and then rapidly quenched in ice water for 60 s to generate monomeric CHP capable of hybridizing with unfolded collagen strands. Tissue sections were incubated with CHP-R overnight at 4 °C and washed three times to remove unbound peptide ^55, 69^, detected in the red channel, was selected instead of CHP-F to minimize interference from the intrinsic green autofluorescence of AF tissue.

AGE accumulation was assessed by UV autofluorescence imaging. Tissue sections were excited at 365 nm, and autofluorescence emission was collected over 420-600 nm, consistent with established AGE detection approaches ^70, 71, 72^. Fluorescent AGEs, including pentosidine, contribute to this UV-induced autofluorescence signal. All sections were imaged using identical exposure and acquisition settings to permit semi-quantitative comparison of fluorescence intensity between non-degenerate and degenerate AF.

### Small- and wide-angle X-ray scattering

Immediately following fatigue testing, IVD specimens (∼800 μm thick) were dehydrated in a desiccator before small- and wide-angle X-ray scattering (SAXS/WAXS) measurements using a SAXSpoint 2.0 system (Anton Paar, Austria). The sample-to-detector distances were 576 mm for SAXS and 113 mm for WAXS. Both measurements used Cu Kα radiation (λ = 1.54 Å) and an Eiger R 1M horizontal detector. For each specimen, three regions along the crack path were analyzed: the crack-tip region immediately adjacent to the crack tip, the mid-crack region located approximately halfway along the crack path, and the crack-end region located within the terminal one-sixth of the crack path. Regional measurement locations are illustrated in Supplementary Fig. 5. Three replicate measurements were acquired from each region, with an acquisition time of 5 min per frame using a 10 × 10 sample stage. Scattering data were processed using SAXSpoint analysis software, including frame averaging, zero-point normalization, beamstop masking, one-dimensional radial integration, and background subtraction. The resulting intensity profiles were fitted using split-Gaussian functions to determine peak positions and full width at half maximum (FWHM). Collagen d-periodicity was calculated from the sixth-order SAXS reflection, while lateral intermolecular spacing and its coherence length were determined from the corresponding WAXS reflection.

### Deep-tissue multiphoton imaging

To enable deep tissue imaging, we used a tissue clearing method based on uDISCO solvent ^73^. The tissue was initially immersed overnight in 4% paraformaldehyde solution for fixation. Dehydration was achieved by sequentially immersing the tissue in ethyl alcohol solutions (30%, 50%, 80%, 90%, 98%, and 100% vol/vol), each for a duration of 2 h. Dichloromethane was then used for a de-lipidation process (30 min). Finally, the tissue was immersed in benzyl alcohol and benzyl benzoate (BABB; 1:2 vol/vol), as well as diphenyl ether (DPE; BABB:DPE = 1:10 vol/vol) overnight. This procedure significantly improved the transparency of tissue samples, allowing for deeper imaging and data acquisition. The tissue was scanned at varying depths using an SP8 multiphoton microscope (Leica, Wetzlar, Germany) with an air objective and a laser excitation wavelength of 910 nm ^74, 75^. The tissue sample remained immersed in the BABB-DPE solution throughout this procedure, thereby preserving its transparency. By scanning collagen structures ranging from superficial layers to deeper formations, we obtained comprehensive images (Supplementary Fig. 4).

### Statistical analysis

Statistical analyses were performed using GraphPad Prism (GraphPad Software, Boston, MA, USA). Data are presented as individual data points with mean ± standard deviation (SD), unless otherwise indicated. Statistical significance was defined as *P* < 0.05.

For comparisons of fracture toughness between non-degenerate and degenerate AF across anatomical loading directions, two-way ordinary ANOVA was performed with tissue condition (non-degenerate versus degenerate; three biological replicates per condition, *n* = 3) and loading direction as factors. Šídák’s multiple-comparisons test was used to evaluate degeneration-associated differences within each loading direction.

To evaluate the effect of proteoglycan depletion on fracture toughness, two-way repeated-measures ANOVA was performed separately for each loading direction. Tissue condition (non-degenerate versus degenerate; three biological replicates per condition, *n* = 3) and enzymatic treatment (untreated versus ChABC-treated) were included as factors. Enzymatic treatment was treated as the repeated-measures factor because untreated and ChABC-treated specimens were obtained from matched tissue samples. Šídák’s multiple-comparisons test was used to assess the effect of ChABC treatment within each tissue condition.

For SAXS/WAXS measurements, two-way repeated-measures ANOVA was performed separately for radial and axial specimens to evaluate the effects of tissue condition (non-degenerate versus degenerate; three biological replicates per condition, *n* = 3) and fatigue loading (intact versus fatigued) on collagen structural parameters, including collagen d-spacing, lateral intermolecular spacing, and WAXS peak full width at half maximum (FWHM). Intact and fatigued specimens were obtained from adjacent regions of the anterior AF within the same disc and were therefore treated as matched samples, with fatigue condition specified as the repeated-measures factor and tissue condition as the between-subject factor. Šídák’s multiple-comparisons test was used for post hoc comparisons of fatigue effects within each tissue condition. For all two-way ANOVA analyses, both main effects and interaction effects were evaluated. An interaction indicates that the effect of one factor depends on the level of the other factor; for example, a significant fatigue × degeneration interaction indicates that the effect of fatigue differs between non-degenerate and degenerate AF. Interaction effects are denoted by “×” throughout the text and are reported where statistically significant.

## Data availability

The source data underlying the findings of this study are provided with the paper. Additional data supporting the findings of this study are available from the corresponding author upon request.

## Code availability

Arduino firmware used to operate the custom fatigue tester is available at https://github.com/xiaoyilan96/Custom-Fatigue-Tester/tree/main.

## Acknowledgements

This work was supported by the Canadian Institutes of Health Research (CIHR; grants PJT-165995 and PJT-203839 to L.H. and J.L.). X.L. was supported by postdoctoral fellowships from the Natural Sciences and Engineering Research Council of Canada (NSERC) and the Fonds de recherche du Québec - Santé (FRQS). N.R.H.S. was supported by an NSERC Canada Graduate Scholarship - Doctoral (CGS-D). A.B. acknowledges financial support from the Fonds de recherche du Québec - Nature et technologies (FRQNT). K.V.P. was supported by an FRQNT Master’s Scholarship. Z. Y. acknowledges support from an FRQNT Doctoral Scholarship. J.L. acknowledges support from the Canada Research Chairs Program.

## Author contributions

X.L.: Conceptualization, Methodology, Investigation, Formal analysis, Data curation, Visualization, Writing - original draft, review & editing. N.R.H.S.: Methodology (SAXS/WAXS), Investigation, Formal analysis, Data curation, Visualization, Writing - original draft, review & editing. T.X.: Methodology (fatigue testing and fatigue-testing system development), Writing - review & editing. H.Z.: Methodology (pure-shear fracture toughness testing), Investigation, Writing - review & editing. K.V.P.: Methodology (fatigue testing), Writing - review & editing. A.B.: Investigation, Formal analysis, Writing - review & editing. Z.Y.: Investigation, Writing - review & editing. L.L.: Methodology (sample collection), Writing - review & editing. Y.X.: Methodology (fatigue), Formal analysis, Writing - review & editing. H.M.T.: Methodology (SAXS/WAXS), Formal analysis, Resources, Writing - review & editing. M.J.H.: Methodology, Resources, Supervision, Writing - review & editing. L.H.: Conceptualization, Methodology, Resources, Supervision, Project administration, Funding acquisition, Writing - review & editing. J.L.: Conceptualization, Methodology, Resources, Supervision, Project administration, Funding acquisition, Writing - review & editing.

## Supplementary information

Supplementary Figs. 1–6.

