## Supplementary Figs. 1-6 and Supplementary Table 1 for "Multiscale fracture resistance of human annulus fibrosus and its deterioration with degeneration"

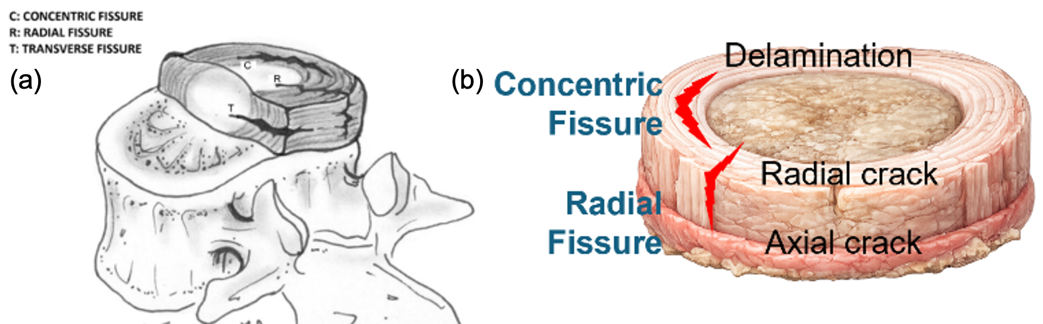

**Supplementary Fig. 1. Annular fissure nomenclature and corresponding experimental crack orientations in the annulus fibrosus (AF).** a Schematic illustration of the three major annular fissure morphologies using standard nomenclature: concentric, radial, and transverse fissures. Concentric fissures propagate between adjacent annular lamellae, approximately parallel to the disc periphery. Radial fissures extend through the annular wall in the radial direction, whereas transverse fissures are oriented approximately horizontally and often occur near the peripheral annulus or vertebral rim attachment. b Correspondence between these clinical fissure morphologies and the crack orientations examined in this study. Delamination cracks represent propagation along interlamellar interfaces and therefore correspond to concentric fissures. Radial and axial cracks both represent through-wall fracture modes related to radial fissure propagation, but differ in the direction of crack advance relative to the local collagen architecture. Transverse fissures were not directly modeled in this study. The clinical fissure schematic in a is reproduced from Fardon et al.^1^, with permission from Elsevier.

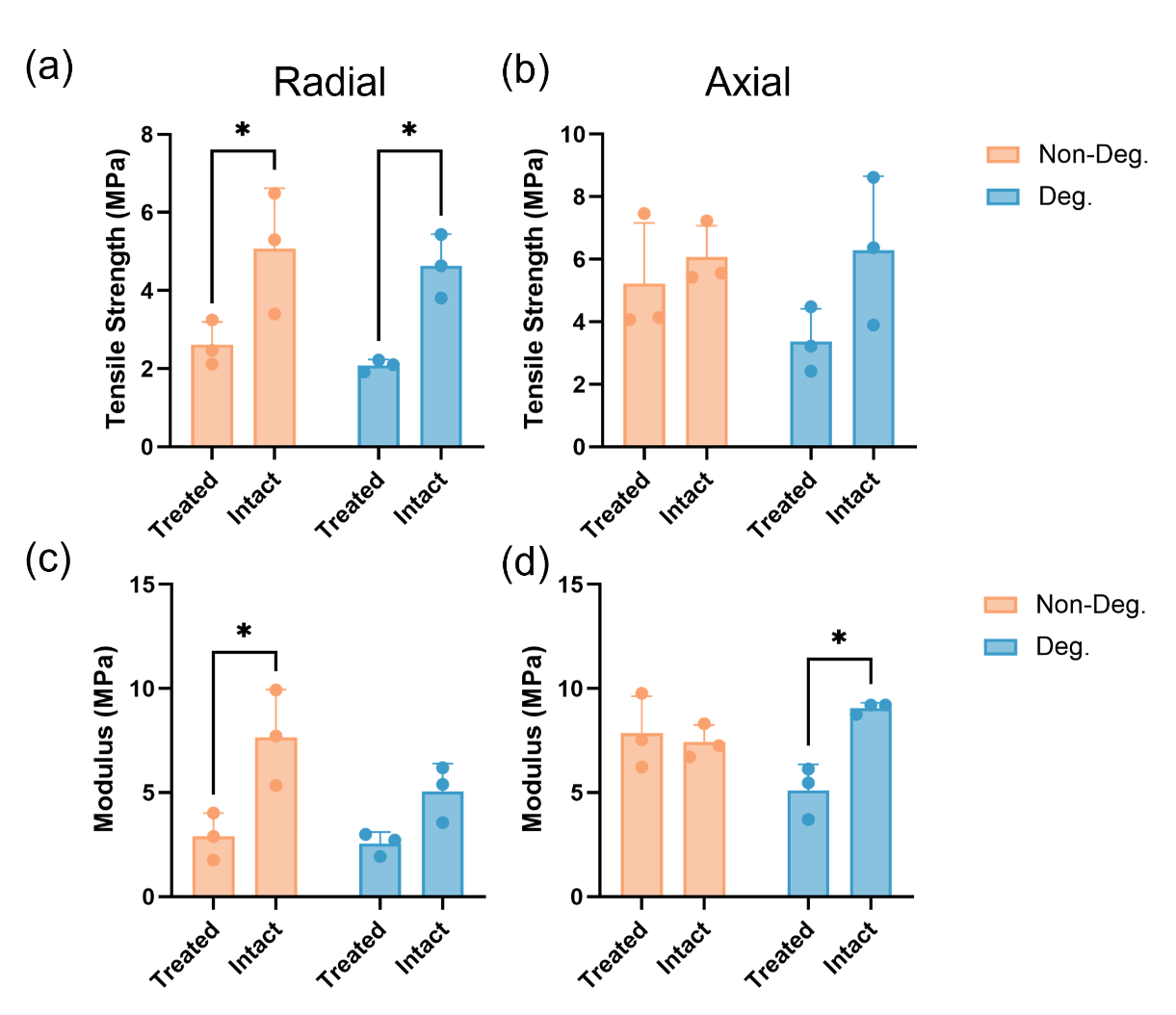

**Supplementary Fig. 2.** **Effect of chondroitinase ABC treatment on tensile properties of the annulus fibrosus.** Tensile properties of non-degenerate and degenerate AF specimens were measured after enzymatic depletion of chondroitin sulfate-associated matrix components using chondroitinase ABC. a,b Tensile strength in the radial and axial directions, respectively. c,d Tensile modulus in the radial and axial directions, respectively. Chondroitinase ABC treatment significantly reduced radial tensile strength in both non-degenerate and degenerate AF, whereas axial tensile strength was not significantly altered. The treatment also reduced tensile modulus in non-degenerate radial specimens and degenerate axial specimens. These results indicate that the contribution of chondroitin sulfate-associated matrix components to bulk tensile behavior depends on both loading orientation and degeneration state. Data are presented as mean ± SD with individual data points shown. Statistical significance is indicated as **P* < 0.05.

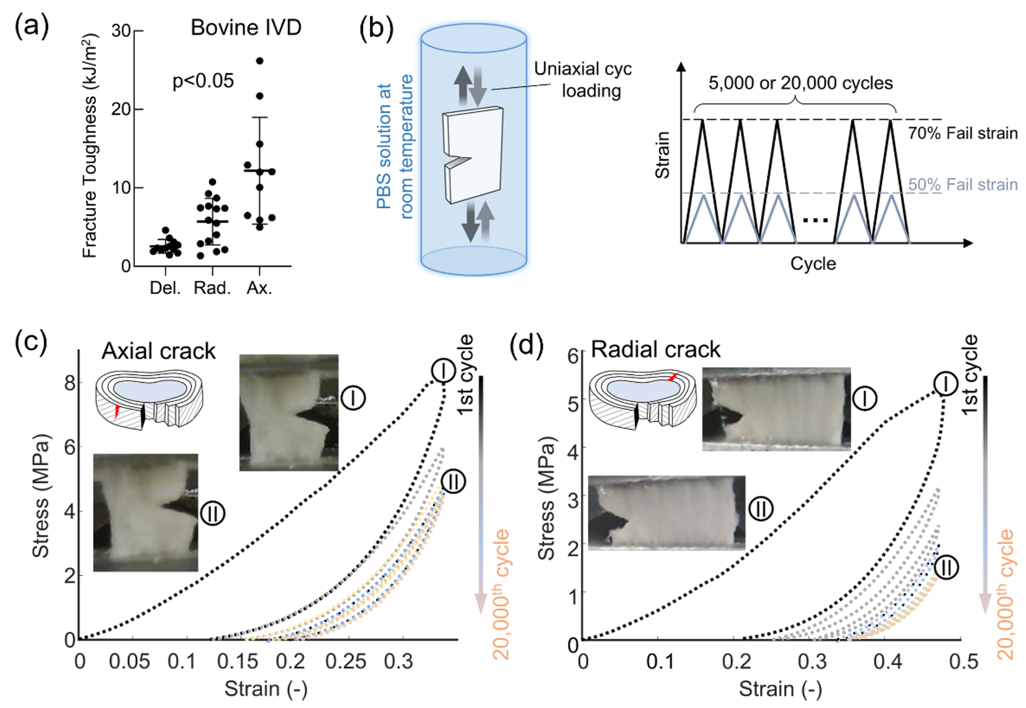

**Supplementary Fig. 3. Mechanical responses of bovine AF.** a Measured fracture toughness for bovine AF. **b** Cyclic loading tests of the edge-notched samples were carried out in PBS solution under 50% and 70% of failure strain for 5,000 or 20,000 cycles. c,d Stress–strain curves of specimens with axial and radial cracks, respectively, during 20,000 loading cycles.

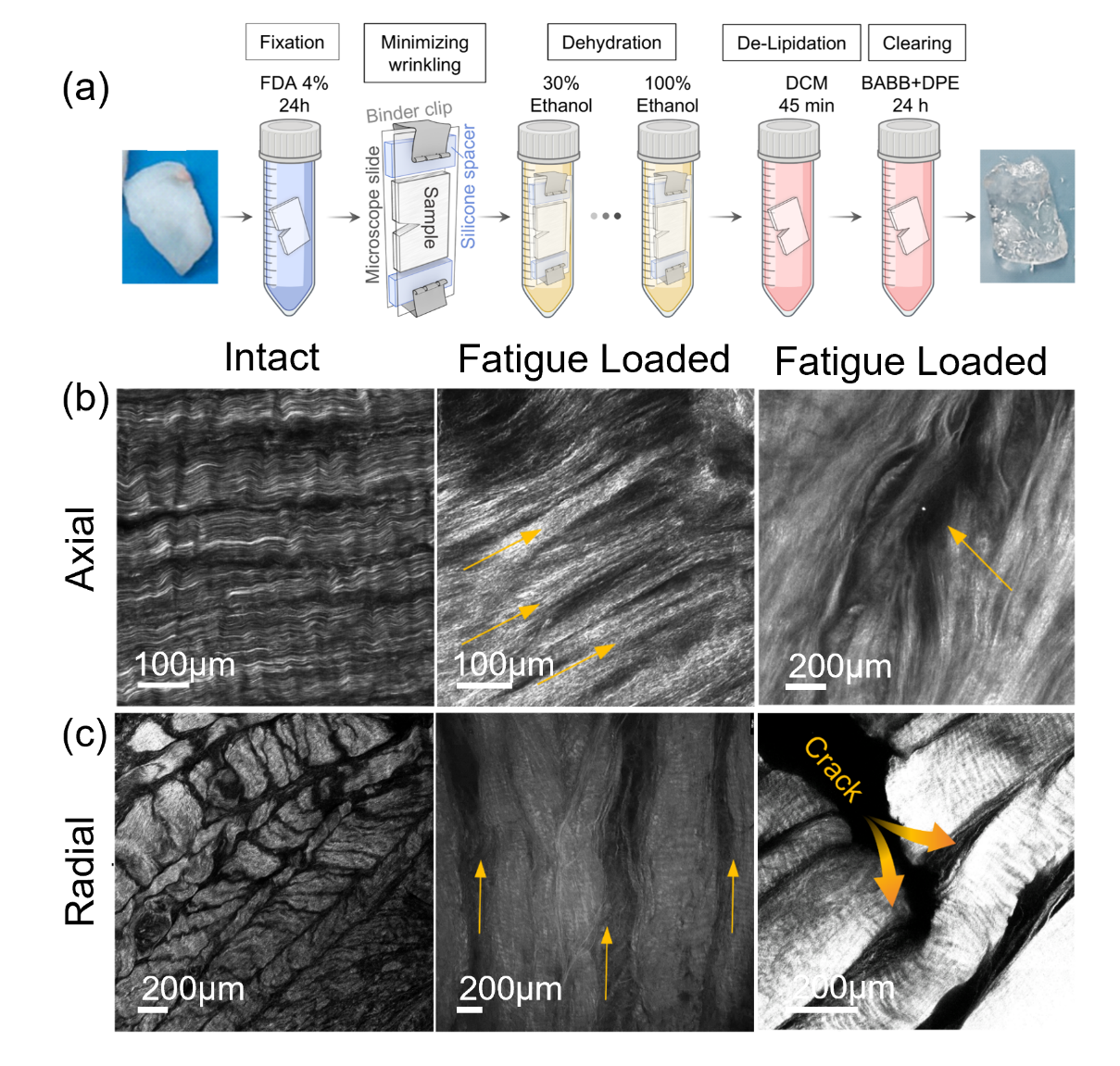

**Supplementary Fig. 4. Microscale characterization of collagen fiber organization and fatigue-associated damage in bovine annulus fibrosus using deep-tissue imaging.** a Schematic of the tissue-processing workflow for uDISCO-based optical clearing, including fixation, minimization of tissue wrinkling, graded ethanol dehydration, delipidation, and refractive-index matching before multiphoton imaging. b Representative multiphoton images of axial AF specimens showing collagen fiber organization in intact tissue and after fatigue loading. Yellow arrows indicate localized changes in fiber alignment and recruitment near the fatigue-damaged region. c Representative images of radial AF specimens showing the multilamellar collagen architecture in intact tissue and fatigue-associated structural changes. Scale bars are indicated in each image.

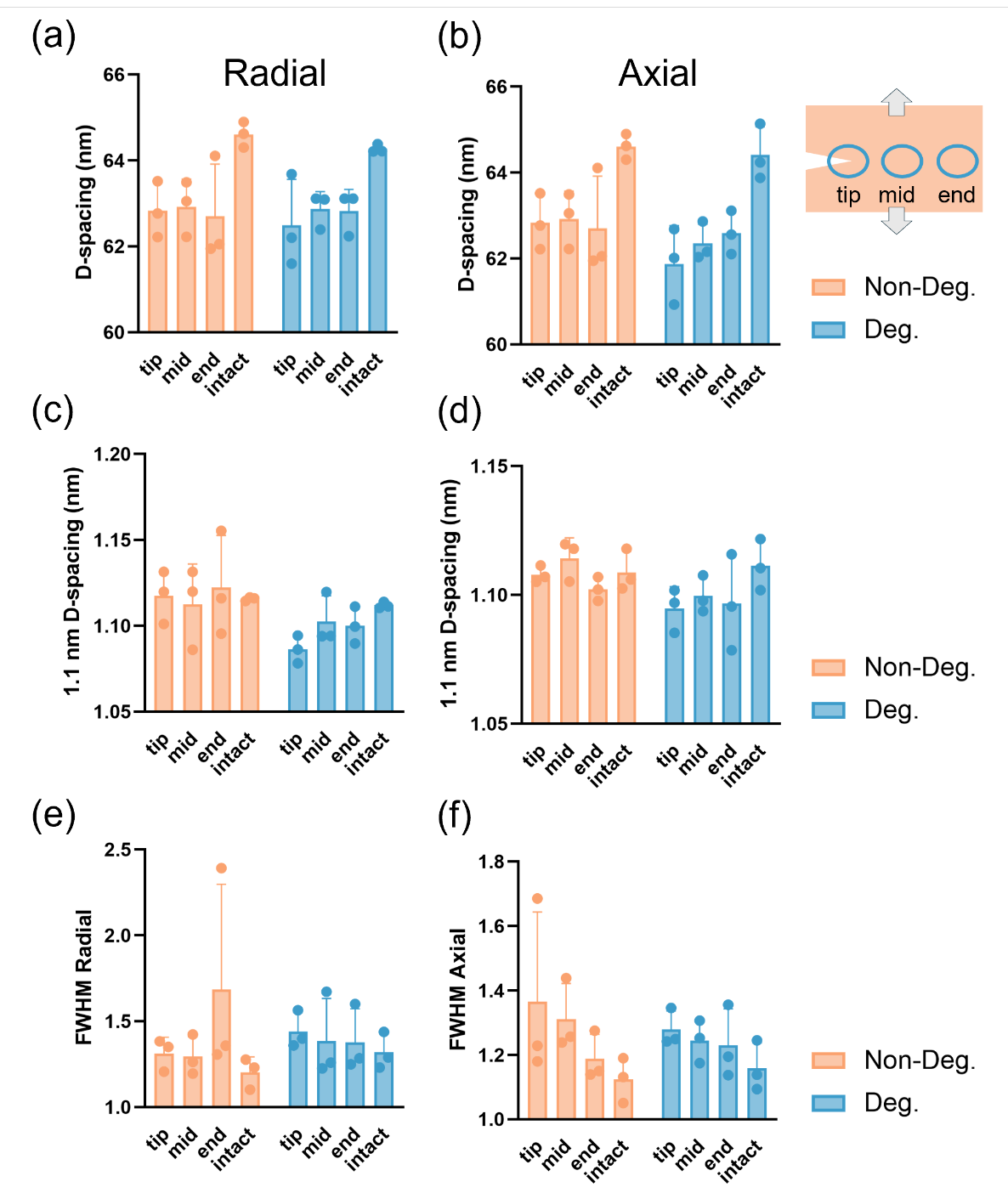

**Supplementary Fig. 5.** **Spatially resolved collagen fibrillar and molecular structural changes along the fatigue crack path.** SAXS/WAXS-derived structural parameters were quantified at defined locations along the crack path, including the crack tip, mid-crack region, and crack end, and compared with intact tissue, as illustrated schematically. a,b Collagen D-period in radial and axial specimens, respectively. c,d Lateral intermolecular spacing associated with the approximately 1.1-nm WAXS peak in radial and axial specimens, respectively. e,f Full width at half maximum (FWHM) of the approximately 1.1-nm WAXS peak in radial and axial specimens, respectively, reflecting the coherence and heterogeneity of lateral intermolecular packing. Measurements are shown separately for non-degenerate and degenerate annulus fibrosus. Together, these spatially resolved measurements characterize persistent fatigue-associated changes in collagen fibrillar periodicity and molecular packing along the crack path. Data are presented as mean ± SD with individual data points shown.

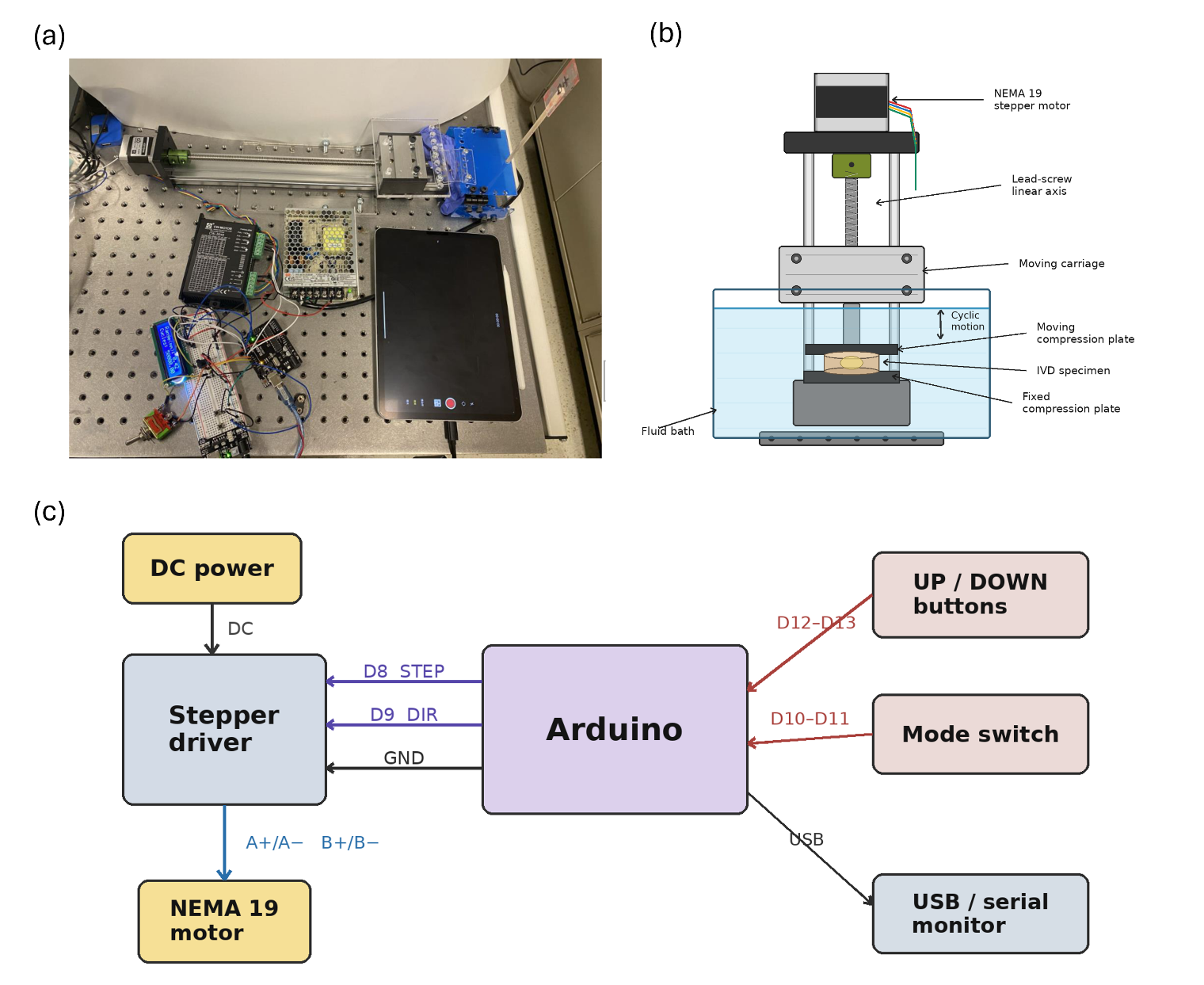

**Supplementary Fig. 6. Custom fatigue tester and electrical control system.** **a** Photograph of the custom-built fatigue tester used for cyclic loading of annulus fibrosus specimens. The system consists of a NEMA 19 stepper motor coupled to a lead-screw linear axis with a movable carriage, a fixed specimen fixture, a stepper motor driver, a DC power supply, and an Arduino-based controller mounted on an optical table. **b** Enlarged view of the specimen-loading region. During testing, the specimen and loading fixtures were immersed in a PBS bath to maintain tissue hydration. **c** Simplified schematic of the electrical control system showing the connections among the stepper motor, motor driver, Arduino controller, manual user controls, LCD display, and USB serial monitor. The Arduino controller operated the tester in displacement-controlled cyclic loading mode using user-defined displacement, velocity, and cycle number. The Arduino firmware used to control the fatigue tester is publicly available at https://github.com/xiaoyilan96/Custom-Fatigue-Tester/tree/main.

**Supplementary Table 1.** Donor characteristics and degeneration grades of human lumbar intervertebral discs.

| **Age/Sex** | **Relevant medical history** | **Cause of Death** | **Disc levels** | **L3/4 Thompson grade** | **L4/5 Thompson grade** | **Classification** |
| --- | --- | --- | --- | --- | --- | --- |
| 72/Male | Kidney stone Osteoarthritis | Cranial Trauma | L5-L4, L4-L3 | V | V | Degenerate |
| 59/Female | Neurodegenerative Disease | Medical assisted death | L5-L4, L4-L3 | IV | IV | Degenerate |
| 63/Male | Blood transfusion | CVA/Stroke | L5-L4, L4-L3 | III-IV | III-IV | Degenerate |
| 21/Male | Cervical contusion/Bruise | Anoxia | L5-L4, L4-L3 | I | I | Non-degenerate |
| 33/Male | Chlamydia Positive 2019, Hepatitis A Positive | Anoxia | L5-L4, L4-L3 | I | I | Non-degenerate |
| 33/Male | Lumbar sprain 2019 | Anoxia | L5-L4, L4-L3 | I | I | Non-degenerate |
